# Identification of 30 transition fibre proteins reveals a complex and dynamic structure with essential roles in ciliogenesis

**DOI:** 10.1101/2023.05.29.542710

**Authors:** Manu Ahmed, Richard Wheeler, Jiří Týč, Shahaan Shafiq, Jack Sunter, Sue Vaughan

## Abstract

Transition fibres are appendages that surround the distal end of mature basal bodies (also called distal appendages) and are essential for ciliogenesis, but only a small number of proteins have been identified and functionally characterised. Here, through genome-wide analysis, we have identified 30 additional transition fibre proteins (TFPs) in the flagellated eukaryote *Trypanosoma brucei* and mapped the arrangement of the molecular components. We discovered TFPs recruited to the basal body pre- and post-initiation of ciliogenesis with differential expression of TFPs at the assembling new flagellum compared to the existing old flagellum. Knockdown by RNAi of 17 TFPs revealed 6 were necessary for ciliogenesis and a further 3 were necessary for normal flagellum length. We identified 9 TFPs that had a detectable orthologue in at least one basal body-forming eukaryotic organism outside of the kinetoplastid parasites. Our work demonstrates that transition fibres are complex and dynamic in their composition throughout the cell cycle which relates to their essential roles in ciliogenesis and length regulation.

## Introduction

Cilia (also called flagella) are highly conserved microtubule-based organelles, which project from the cell surface and play essential roles in signalling and/or motility in a range of eukaryotic organisms (Moran et al., 2014). They are assembled from microtubule-based basal bodies (also called centrioles), which are composed of nine radially arranged microtubule triplets. Basal bodies and centrioles often exist as a connected pair and duplicate in a cell cycle-dependent manner with each daughter cell inheriting an older (mature) basal body and a younger (immature) basal body. Only a mature basal body can assemble a cilium/flagellum and contain additional appendages called transition fibres (TFs) (called distal appendages in mammalian cells), which are a set of nine blade-like structures that radiate out from the triplet microtubules at the distal end of the mature basal body.

In mammalian cells, assembly of a cilium occurs during G0 following mitotic exit. During ciliogenesis, the mature basal body docks to the plasma membrane via distal appendages that anchors the cilium as it extends out from the cell (Anderson, 1972; O’Hara, 1970). To date at least 10 proteins have been identified at transition fibres in mammalian cells – C2CD3, CEP83, CEP90, OFD1, CEP89, SCLT1, CEP164, ANKRD26, CEP19 and FBF1 with important functions in membrane docking and modulating ciliogenesis (Bowler et al., 2019; Kurtulmus et al., 2018; Tanos et al., 2013; Yang et al., 2018; Kumar et al., 2021). Defective cilia are linked to a class of human diseases known as ciliopathies. Mutations in transition fibre proteins are linked to a number of syndromic ciliopathies, demonstrating the importance of distal appendages for cilium formation and function (Failler et al., 2014; Chaki et al., 2012).

Trypanosomes are an excellent system to study ciliogenesis as they possess a basal body pair and a single flagellum that undergoes cell cycle duplication, maturation and anchoring in a similar manner to mammalian basal bodies. The flagellum also exhibits the same canonical features with a transition zone, transition fibres and a 9 + 2 axoneme that undergo cell cycle-regulated assembly (Sherwin and Gull, 1989; Wheeler et al., 2019; Hodges et al., 2010). The genome of trypanosomes encodes many known conserved centriole and cilia genes including many genes involved in ciliopathy diseases of humans (Broadhead et al., 2006; Hodges et al., 2010; Barker et al., 2014). Therefore, insights gained from trypanosomes are likely to apply to other eukaryotic organisms. The main advantages of this model organism for flagellum biogenesis studies are the ability to follow the assembly and duplication of basal bodies and unequivocally identify old flagella and assembling new flagella in dividing cells. A G1 trypanosome cell has one mature and probasal body pair located at the proximal end of a single flagellum (Fig 1A; G1). This flagellum does not disassemble and a new flagellum grows alongside the existing flagellum in a precise position in preparation for division (Fig 1A; S). Assembly of the new flagellum begins with basal body maturation, acquisition of transition fibres and docking of the newly mature basal body to the flagellar pocket occurring near the G1/S-phase boundary. New pro-basal bodies form during S-phase next to each mature basal body and the new flagellum continues growth throughout the rest of the cell cycle (Fig 1A; S) (Wheeler et al., 2019; Sherwin and Gull, 1989; Gluenz et al., 2011).

**Figure 1:**
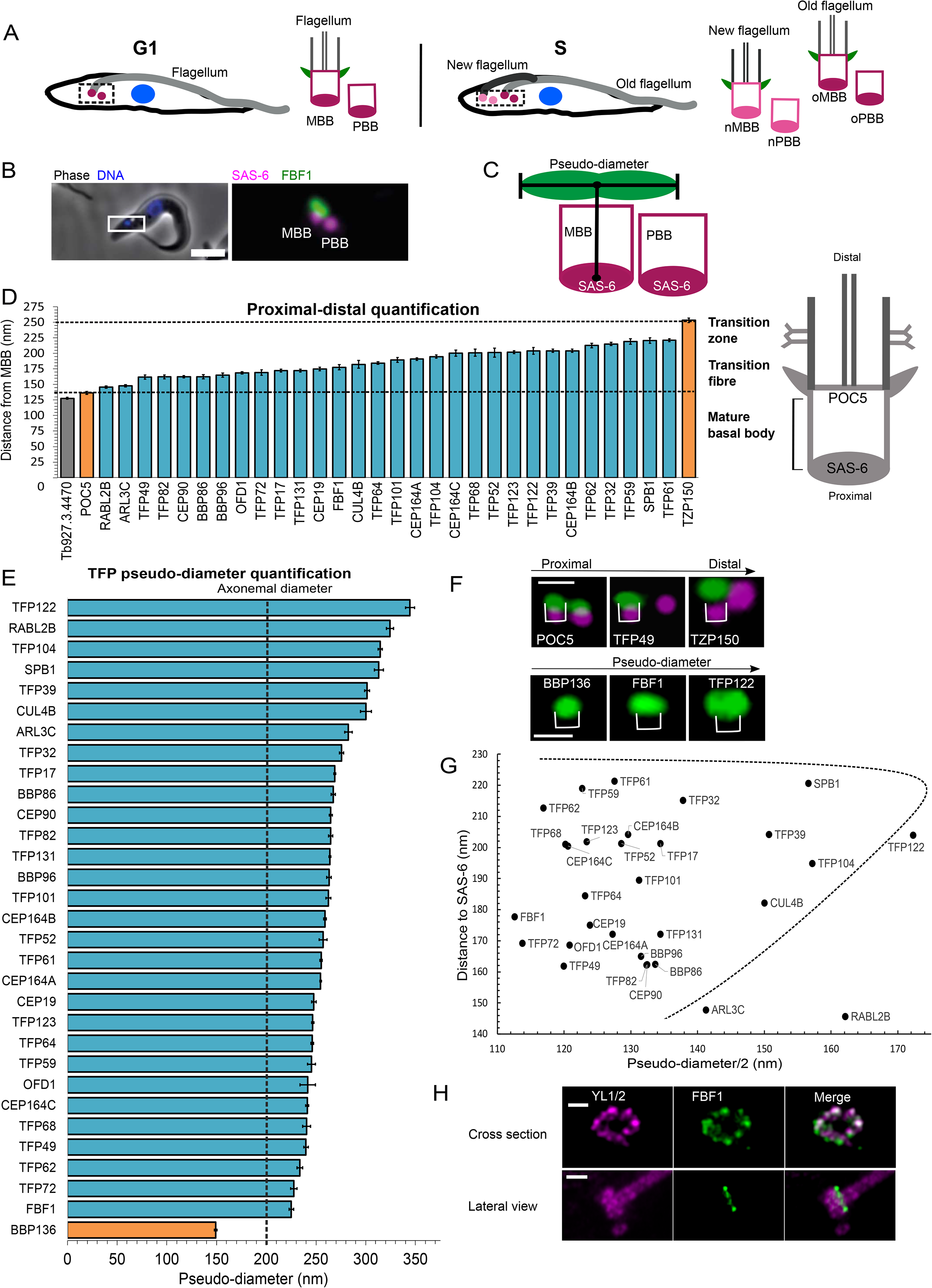
Spatial organisation of candidate transition fibre proteins. A: Cartoon representing trypanosome cell, basal bodies (magenta), transition fibres (green) and axoneme (grey). G1 cell (left) represented with a single flagellum (light grey) and basal body pair (magenta). S-phase cell with new flagellum (dark grey) and old flagellum (light grey) each with a corresponding basal body pair at the new (pink; nMBB, nPBB) and old flagellum (magenta; oMBB, oPBB). MBB = mature basal body, PBB = pro-basal body, o=old, n=new. B: TF protein endogenously tagged with FBF1:mNG and SAS-6:mScarlet. FBF1 is only located on MBB; C: cartoon illustration of distance and pseudo-diameter measurements carried out in D and E – see materials & methods; D: Range of measurements of candidate TFPs ordered by their distance from SAS-6 (nm). Controls – POC5 for MBB distal end. Candidates with a shorter distance were excluded (grey). TZP150 for transition zone. E: Measurements ordered by pseudo-diameter (nm). Basal body control (BBP136) shown in orange, theoretical axonemal diameter indicated by the grey line at 200 nm. F: Examples of TFP localisations based on their proximal-distal location (top) and pseudo-diameter (below). Scale bar = 500 µm. G: Model representing the approximate location of each transition fibre protein group based on diameter and distance from basal body (SAS6). TZ – transition zone, TF – transition fibre; H: Expansion microscopy (ExM) of FBF1::TY co-localised with anti- tubulin antibody YL1/2. Scale bar = 500 nm.

Scanning transmission electron tomography confirmed the canonical nine blades of transition fibres docked to the flagellar pocket in trypanosomes (Trépout et al., 2018). In addition, two widely conserved proteins - CEP164C and Retinitis Pigmentosa-2 (RP-2) have been localised at transition fibres (Stephan et al., 2007; Harmer et al., 2017; Atkins et al., 2021) and a kinetoplastid-specific TFK1 localised to transition fibres with CEP164C and RP-2 (Ramanantsalama et al., 2022). Overall in molecular terms, transition fibres have broadly been considered a simple structure comprising only a handful of proteins. Using a genome- wide protein localisation resource (Billington et al., 2023), our study has identified 30 transition fibre proteins, mapping their position, timing of incorporation and carried out functional studies using RNAi. We conclude transition fibres are complex and dynamic structures comprising a core set of evolutionarily conserved components that are essential for assembly of a eukaryotic cilium or flagellum.

## Results

### Identification of 30 transition fibre proteins

A genome-wide protein localisation study identified ∼307 putative basal body proteins based on the localisation to one or two small foci at the proximal end of the single trypanosome flagellum (Billington et al., 2023). Since transition fibres are only on the mature basal body, proteins with a clear single focus at the proximal end of a flagellum were taken forward for further study to search for novel transition fibre proteins. Localisation to near the mature basal body was confirmed by expressing the putative transition fibre proteins with an mNeonGreen (mNG) tag in a cell line expressing the basal body marker SAS-6 endogenously tagged with mScarlet (Nakazawa et al., 2007; Fong et al., 2014). In total, 31 candidates with a focus at the mature basal body were taken forward. An example is the ortholog of characterised mammalian TF protein FBF1 which is located on the mature basal body only, as expected (Fig 1B) (Tanos et al., 2013). All hypothetical proteins were named transition fibre protein (TFP) followed by their theoretical molecular weight.

Transition fibres are located at the distal end of a mature basal body and surround the basal body barrel, so we reasoned that if a candidate was likely to represent a TFP candidate then the fluorescence signal would be expected to be distal to the location of SAS-6, which sits at the proximal base of the basal body barrel. In addition, TFP signal might be expected to have a wider diameter as they are located around the outside of the basal body barrel (Fig 1C). We refer to the length of the signal from a lateral view as the pseudo- diameter (Fig 1C). Quantification of distance from SAS-6 and pseudo-diameter of candidate signal was carried out using an automated FIJI script and measurements were made on up to 2000 methanol-fixed cytoskeletons per candidate and was plotted for each putative TFP (Fig 1D, E). Measurements were made on live cells for TFPs that were not observed following detergent treatment with 1% Nonidet P-40 (ARL3C, RABL2B and TFP122). Firstly, all candidates that were both distal to SAS-6 and POC5 (Fig 1F) (POC5 is a known distal end basal body protein, but not a TFP) (Azimzadeh et al., 2009), but more proximal than the transition zone protein TZP150 (Dean et al., 2016) (Fig 1D, F) were taken forward. These criteria removed one candidate (Fig 1D; Tb927.3.4470). Secondly, all candidates with a pseudo-diameter greater than BBP136 (a bona fide mature basal body protein) (Dang et al., 2017) (Fig 1E, F) were taken forward for further study (Fig 1E) and examples (Fig 1F). This latter criteria did not remove any further candidates. Known transition fibre orthologues in trypanosomes identified by this method - CEP164A, CEP164B, CEP164C (Hodges et al., 2010) and OFD-1 (Andre et al., 2014) -all fit within the expected region, demonstrating that this method can successfully identify transition fibre proteins (Fig 1D, E). These two measurements resulted in a total of 30 TFPs being taken forward for further study (Supplemental Fig 1). Finally, to further confirm that candidates are located at TFs, a cell line was generated expressing each TFP candidate tagged with mNG and mScarlet tagged CEP164A. For those proteins we were unable to successfully generate cell lines, co-localisation was carried with an antibody to Retinitis Pigmentosa-2 (YL1/2) which localises to the TFs in Trypanosomes (Stephan et al., 2007; Atkins et al., 2021)(Supplemental Fig 2). Fig 1G indicates the approximate location of each TFP based on distance and pseudo-diameter measurements.

There was a range of pseudo-diameter measurements consistent with there being TFPs located along the length of each transition fibre blade (Fig 1E). TFP122, RABL2B, TFP104, SPB1, TFP39, CUL4B exhibited the widest pseudo-diameter (300 - 345 nm) (Fig. 1E, G) and the rest a narrower pseudo-diameter (225 - 283nm) (Fig 1E, G). To compare to the ultrastructural dimensions of TFs to our light microscopy measurements, we analysed the distance between the most outer most regions of TFs by longitudinal thin section transmission electron microscopy (Supplemental Fig 3). TFs showed a mean diameter of 354.3±10.8 nm (n=20) by EM, which is a similar range to wider proteins in our light microscopy datasets (Supplemental Fig 3). Finally, expansion microscopy (Gorilak et al., 2021; Kalichava and Ochsenreiter, 2021) was also carried out on FBF1 using an endogenous TY epitope tag. This demonstrated an approximate 9-fold transition fibre arrangement around the microtubules of the basal body, providing further confirmation of these proteins being associated with the transition fibre area of trypanosome cells (Fig 1H). In summary, this screen has identified 30 TFPs in trypanosomes, significantly extending our molecular map of transition fibres (Supplemental Table 1).

### Evolutionary conservation of TFPs

To analyse the evolutionary conservation of the proteins identified in this screen, reciprocal best BLAST and Orthofinder (Emms and Kelly, 2015) were used to identify orthologues in organisms whose ability to assemble basal bodies is known (Carvalho-Santos et al., 2011a). Our data showed a correlation with the ability to form basal bodies and presence of orthologues of *T. brucei* transition fibre protein-coding genes within the genome, indicating a likely specific requirement in ciliogenesis (Fig 2A; light grey). In this set we included 4 organisms (*Caenorhabditis elegans, Drosophila melanogaster, Plasmodium falciparum, Trichomonas vaginalis*) that have basal bodies, but do not appear to have structures resembling transition fibres by electron microscopy (Carvalho-Santos et al., 2011a). However, both *T. vaginalis* and *D. melanogaster* have orthologues of CEP164 and recently, CEP164, FBF1 and CEP89 were experimentally characterised in *D. melanogaster* located to the distal end of the basal bodies that suggests additional functions for these proteins (Hou et al., 2023). 9 TFPs identified in our screen had a detectable ortholog in at least one basal body-forming eukaryotic organism outside of kinetoplastid parasites (RABL2B, CEP164, CEP19, OFD1, CEP90, FBF1, TFP101, TFP39 and TFP82), with the most widely conserved being CEP164, CEP19 and RABL2B. The remaining TFPs were only detected in other kinetoplastids. From this analysis we can concluded that whilst transition fibres appear to display many structural and functional similarities across ciliated cells, TFPs are either not well conserved across the wide range of ciliated cells in eukaryotic organisms or do not have a well conserved primary sequence.

**Figure 2:**
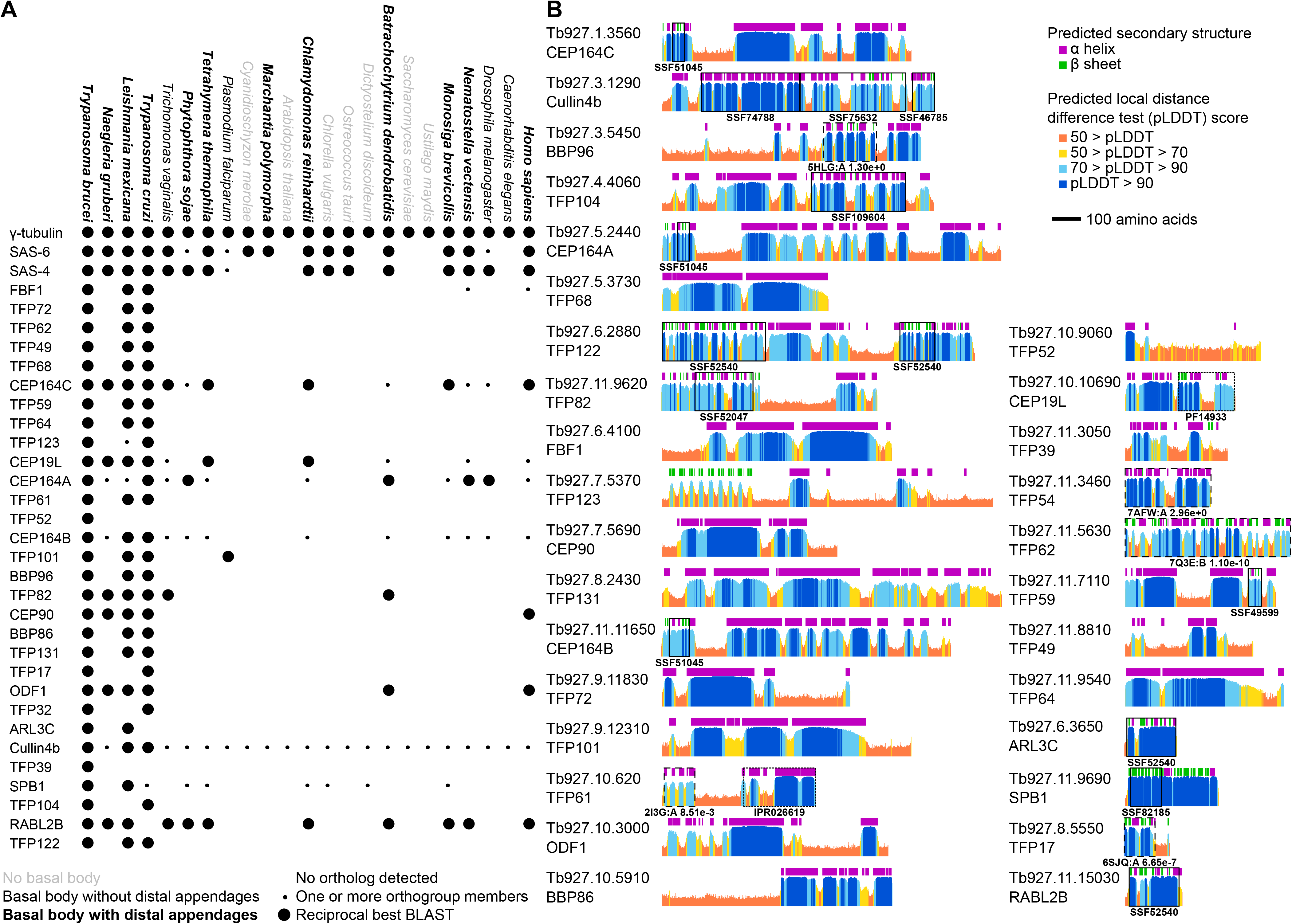
Evolutionary conservation profile and protein structure prediction of T. brucei TFPs. A: Presence or absence of orthologues of *T. brucei* TFPs across diverse eukaryotes, in comparison to control tubulin nucleation (γ-tubulin) and basal body proteins (SAS-6, SAS-4). Large circles indicate presence of an ortholog detectable by reciprocal best BLAST, small circles indicate one or more orthogroup members but not a reciprocal best BLAST. Species fonts indicate their ability to form basal bodies and distal appendages, as previously described (Carvalho-Santos et al., 2011a); B: Alphafold2 structure predictions of all *T. brucei* TFPs. As a large majority of predictions showed extended alpha helices or no predicted structure showing the structures is often not highly informative. Instead, we summarise the structures as per residue plots of predicted local distance difference test (pLDDT) and predicted secondary structure, using the dictionary of secondary structure program (DSSP) on the predicted structure. Protein domains predicted by sequence similarity are shown with boxes: solid boxes for superfamily domains (accessions starting SS), dotted boxes for other database predictions (reserved for when no superfamily domain hit was in that protein region). Dashed boxes show globular regions where we carried out structural similarity comparisons to experimentally determined structures using FoldSeek.

Fewer than half of *T. brucei* TFPs have predicted protein domains detectable by sequence (superfamily database (Wilson et al., 2009)) and these rarely span a large portion of the protein sequence. Therefore, we explored the AlphaFold2 predicted structure (Jumper et al., 2021b), using lineage-optimised input multiple sequence alignment (Wheeler, 2021), to gain insight into potential function (Fig 2B). Many are predicted to be extremely alpha helix rich. FBF1, TFP68, TFP64, TFP101, TFP131 are predicted to be essentially entirely alpha helical. Most have long helical regions, sometimes also with small globular domains such as the CEP164 family with an N terminal WW domain (SSF51045) and long C terminal alpha helix domains. These have the potential to be physically large; with a pitch of 5.4 Å and 3.6 amino acids per turn, 1000 amino acids could form a 150 nm alpha helical arm. For protein regions with a predicted globular structure but no corresponding superfamily domain, we searched for the protein with the most similar experimentally-determined structure in Protein Data Bank (PDB) using FoldSeek (Kempen et al., 2022). TFP17 has a similar predicted structure to *T. brucei* BILBO1 (e value 6.65x10^-7^ vs PDB 6SJQ) (Florimond et al., 2015), a major component of the flagellar pocket, which the trypanosome basal bodies dock onto for ciliogenesis. TFP62 has a similar predicted structure to mouse *inturned* (e value 1.10x10^-10^ vs PDB 7Q3E chain B), part of the ciliogenesis-associated CPLANE complex (Langousis et al., 2022). Only one protein had any similarity to the predicted structure of TFP32 (e value 2.96x10^0^ vs 7AFW chain A). With such a marginal e value, this should likely be discounted–however it is notable that this hit is β-catenin, which has known cilia-associated functions. The C terminus of TFP122 has a domain with predicted structure similar to pleckstrin homology domain, typically involved in phosphatidylinositol binding. Consistent with its wide pseudo-diameter, TFP122 may therefore be involved in membrane interaction. From this data we can conclude that whilst there appears to be a core set of TFPs that are at least conserved between humans and trypanosomes, the nature of TFPs means that they may diverge in sequence too greatly to identify orthologs in distant species by sequence methods alone across eukaryotic organisms.

### Transition fibre proteins are recruited pre- and post-initiation of ciliogenesis

The acquisition of transition fibres by a mature basal body is a critical step in docking and ciliogenesis (Wei et al., 2015; Tanos et al., 2013; Shmidt et al., 2012). Docking of a new mature basal body and growth of a new flagellum narrowly precedes the formation of two new pro-basal bodies in trypanosomes (Fig 1A) (Gluenz et al., 2011; Wheeler et al., 2019). This feature allows us to discover if TFPs arrive as docking of the mature basal body occurs or after initiation of ciliogenesis. SAS-6 is one of the earliest proteins to arrive at the start of new pro-basal body assembly (Hu et al., 2015). Therefore, we used cell lines expressing SAS- 6 tagged with mScarlet with each mNG tagged TF protein as a temporal marker of pro-basal body formation. This allows us to determine if a TFP was recruited to the mature basal body of the new flagellum prior to pro-basal body assembly (before initiation of ciliogenesis) or are recruited to the new mature basal body post basal body duplication (after initiation of ciliogenesis).

FBF1 had a single wide signal overlying the mature basal body (MBB) in cells with 2 SAS-6 foci (Fig 3A; G1 cells). In a proportion of 2 SAS-6 foci cells there was a V-shaped signal representing the transition fibres of two separate, but closely positioned mature basal bodies (Fig 3A; G1/S). This demonstrates early recruitment to the new mature basal body before basal body duplication and, therefore is being recruited prior to ciliogenesis. A total of 15 TFPs were recruited to the new mature basal body prior to ciliogenesis and represent early markers of transition fibre acquisition as docking occurs (FBF1, TFP104, TFP101, TFP61, TFP39, TFP59, TFP49, TFP64, BBP96, BBP86, TFP123, TFP131, TFP17, TFP72, TFP82) (Fig 2A).

**Figure 3:**
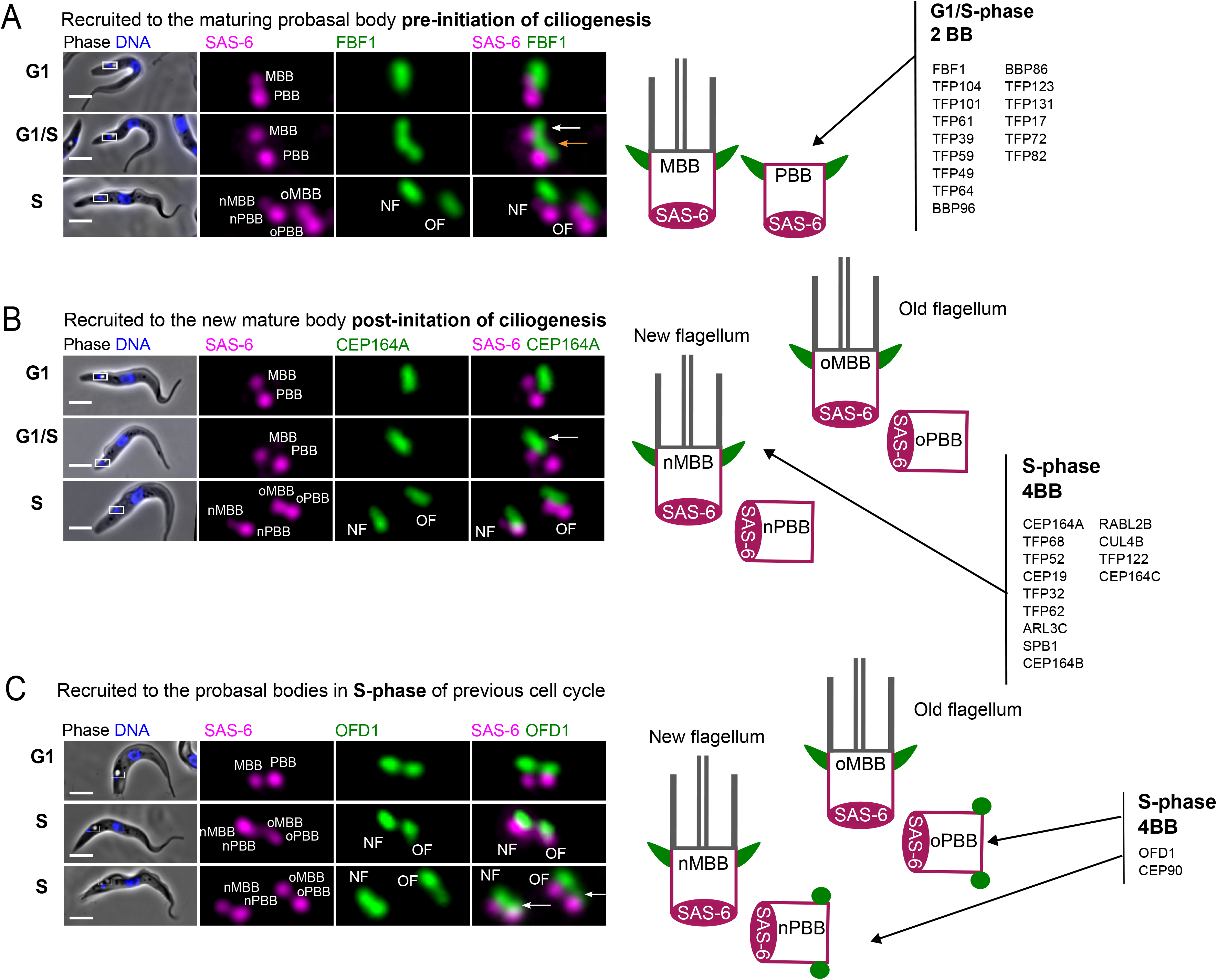
Transition fibre protein recruitment pre- and post-initiation of ciliogenesis. A: Example of a TFP recruited prior to basal body duplication. Images of a live cell expressing endogenously tagged FBF1:mNG (green) and SAS-6::mSc (magenta) from G1 to S-phase (left). At G1/S an FBF1 foci can be observed on the existing mature basal body (white arrow) and a second FBF1 foci can be observed on the maturing body prior to basal body duplication (orange arrow). Cartoon representing localisation of TFPs recruited before basal body duplication in trypanosomes (right); B: Example of a TFP recruited after basal body duplication. Images of a live cell expressing CEP164A::mNG (green) and SAS-6::mSc (magenta) from G1 to S-phase (left). At G1/S the CEP164A foci can only be observed on the existing mature basal body (white arrow) and a second CEP164A foci is not observed on the maturing body prior to basal body duplication (4 X SAS-6). C: Example of a TFP recruited to probasal basal bodies in S-phase of the previous cell cycle. Images of a live cell expressing OFD1::mNG (green) and SAS-6::mSc (magenta) from G1 to S-phase (left). Post basal body duplication, OFD1 foci can be observed on the mature basal bodies (S; middle, white arrows) and subsequently observed arriving on both probasal bodies (S; lower). Index: oPBB – old pro-basal body, nMBB – new mature basal body, nPBB – new pro-basal body.

In contrast, CEP164A is an example of later recruitment to a mature basal body transition fibres after pro-basal body assembly and, therefore, is being recruited after initiation of ciliogenesis. A second signal was not detectable until there were 4 SAS-6 foci (Fig 3B; S - 4 SAS-6 foci). In total 13 TFPs were recruited after pro-basal body assembly (CEP164A, TFP68, TFP52, CEP19, TFP32, TFP62, ARL3C, SPB1, CEP164B, RABL2B, CUL4B, TFP122 and CEP164C) (Fig 3B).

Finally, CEP90 and OFD1 did not fit into either pattern above but were located on the mature and pro-basal body in G1 (Fig. 3C; G1). In S-phase cells (4 SAS-6 foci), OFD1 and CEP90 were observed on the mature basal body (Fig. 3C; S middle) and were subsequently recruited to the pro-basal bodies later in the cell cycle (Fig 3C; S lower panel). The data suggest OFD1 and CEP90 localise to the pro-basal bodies in S-phase of the previous cell cycle, immediately after their assembly and before they start maturation in the next cell cycle. Therefore OFD1 and CEP90 are the earliest recruited TFPs. Taken together, the data is consistent with transition fibres having a hierarchical recruitment process of TFPs as docking an initiation of ciliogenesis is taking place.

### Differential localisation patterns of TFP122, RABL2B, TFP104 and TFP39 reveal differences between flagella of different ages

An important advantage of working with *T. brucei* cells is the ability to determine the precise ‘age’ of the two flagella. During the cell cycle, the old flagellum remains assembled, and a new flagellum grows alongside in a consistent relative position (Fig 1A). In our previous work we showed that CEP164C was only recruited to the transition fibres of the old flagellum. Thus, recruitment only occurred in the third cell cycle after the mature basal body was originally assembled as a pro-basal body. Furthermore, functional studies revealed CEP164C to be essential for regulation of flagella length where it is proposed to be part of a locking mechanism to prevent further growth of the old flagellum whilst the new flagellum assembled during the cell cycle (Bertiaux et al., 2018; Bertiaux and Bastin, 2020; Atkins et al., 2021). Analysis of images of all 30 tagged TFPs revealed that whilst most appeared to have an equal intensity at the base of old and new flagella, TFP122, RABL2B, TFP104, TFP39 exhibited intensity differences between the two flagella.

In RABL2B, TFP39, TFP104 and TFP122 expressing cells there was a significantly higher signal intensity on the transition fibres of the assembled old flagellum (OF) relative to the assembling new flagellum (NF) in dividing cells (Figure 4A-D, F), similar to CEP164C. This differential level of expression remained throughout the cell cycle. We can conclude that these proteins are upregulated following cytokinesis when the flagellum has reached the correct length and may play a role in preventing further growth of the old flagellum. Finally, TFP72 was identified with a significantly higher signal intensity on the TFs of the assembling new flagellum (NF) through the cell cycle (Fig 4E, G), suggesting that the protein is upregulated in the new flagellum during assembly and is then down regulated once assembly is complete at the end of the cell cycle.

**Figure 4:**
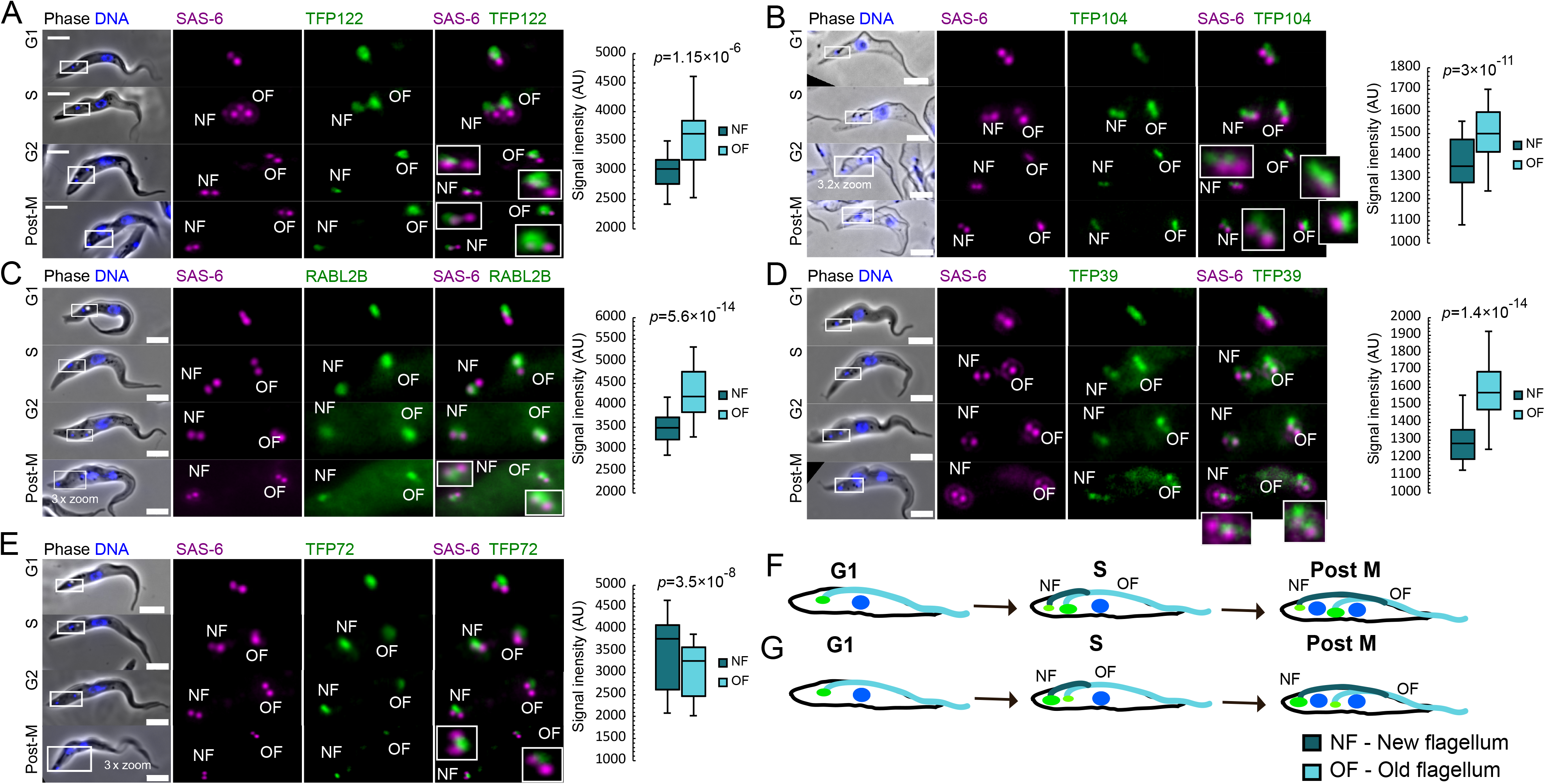
Differential localisation of TFPs at the new and old flagellum A: Live cell images showing the localisation of TFPs throughout the cell cycle (left) and quantification of signal intensity at the old and new flagellum (right) of TFP122 (A), TFP104 (B), RABL2B (C), TFP39 (D) and TFP72 (E). Cell cycle stages are approximate indicated as G1 (2 basal bodies, 1 kinetoplast, 1 nucleus), S (4 basal bodies, 1 kinetoplast, 1 nucleus), G2 (4 basal bodies, 2 kinetoplast, 1 nucleus) and post-mitotic (4 basal bodies, 2 kinetoplast, 2 nuclei). Scale bar = 5µm. NF – new flagellum, OF – old flagellum. Inset represents a 5x zoom unless indicated otherwise in figure. F: Cartoon illustrating increased signal intensity at the TFs at the base of the old flagellum (OF) through the trypanosome cell cycle. G: Cartoon illustrating increased signal intensity at the TFs at the base of the new flagellum (NF) through the trypanosome cell cycle.

### Functional screen reveals essential role for TFPs in flagellum assembly and length regulation

In order to establish if TFPs were essential for flagellum assembly or flagellum length regulation a total of 17 TFPs were selected for functional studies based on their bioinformatic and/or expression profiles using the well characterised stable inducible RNAi knockdown system in trypanosomes (Bellofatto and Palenchar, 2008). The RNAi constructs for the 17 TFPs were integrated into a cell line expressing the target TFP endogenously tagged with mNG, for use as a reporter of knockdown, and CEP164A endogenously tagged with mScarlet to establish if knockdown of the TFP perturbed flagellum growth and/or CEP164A localisation. Once stable cell lines were generated and then individual RNAi cell lines were induced with doxycycline and compared with the uninduced cell population. Inspection of mNG fluorescence by microscopy was used to confirm knockdown of target TFPs from 24hrs post-induction of RNAi. Cells were labelled with monoclonal antibodies (mAb25 - Dacheux et al., 2012) to detect the axoneme of the flagellum and BBA4 (Woodward et al., 1995) to detect basal bodies. Length measurements were taken of new and old flagella in cells with 2 flagella (Post M) and in cells with one flagella (G1) to assay defects in flagellum assembly and length regulation.

6 TFPs were essential for new flagellum assembly, but not for basal body duplication (Fig 5A-F). In CEP90 and TFP68 depleted cells, new flagellum growth was severely perturbed and cell growth was reduced (Fig 5A; left panel). 85% of CEP90 RNAi dividing cells had no new flagella or only a very short new flagellum (mean average 3.1μm) by 48 h post- induction with 61% having no visible flagellum. In addition, CEP164A failed to localise to transition fibres, but basal body pairs were still present (Fig 5A). In TFP68 induced cells, new flagellum growth was also perturbed with very short new flagella (Fig 5B) and there was a wide variation in flagellum length. Furthermore, CEP164A also failed to localise to the transition fibres, but basal bodies were still present in cells, even in if there was no visible flagellum. This indicates that basal body duplication and positioning was unaffected even though recruitment of at least CEP164 is perturbed in both CEP90 and TFP68 (Fig. 5A, B).

**Figure 5:**
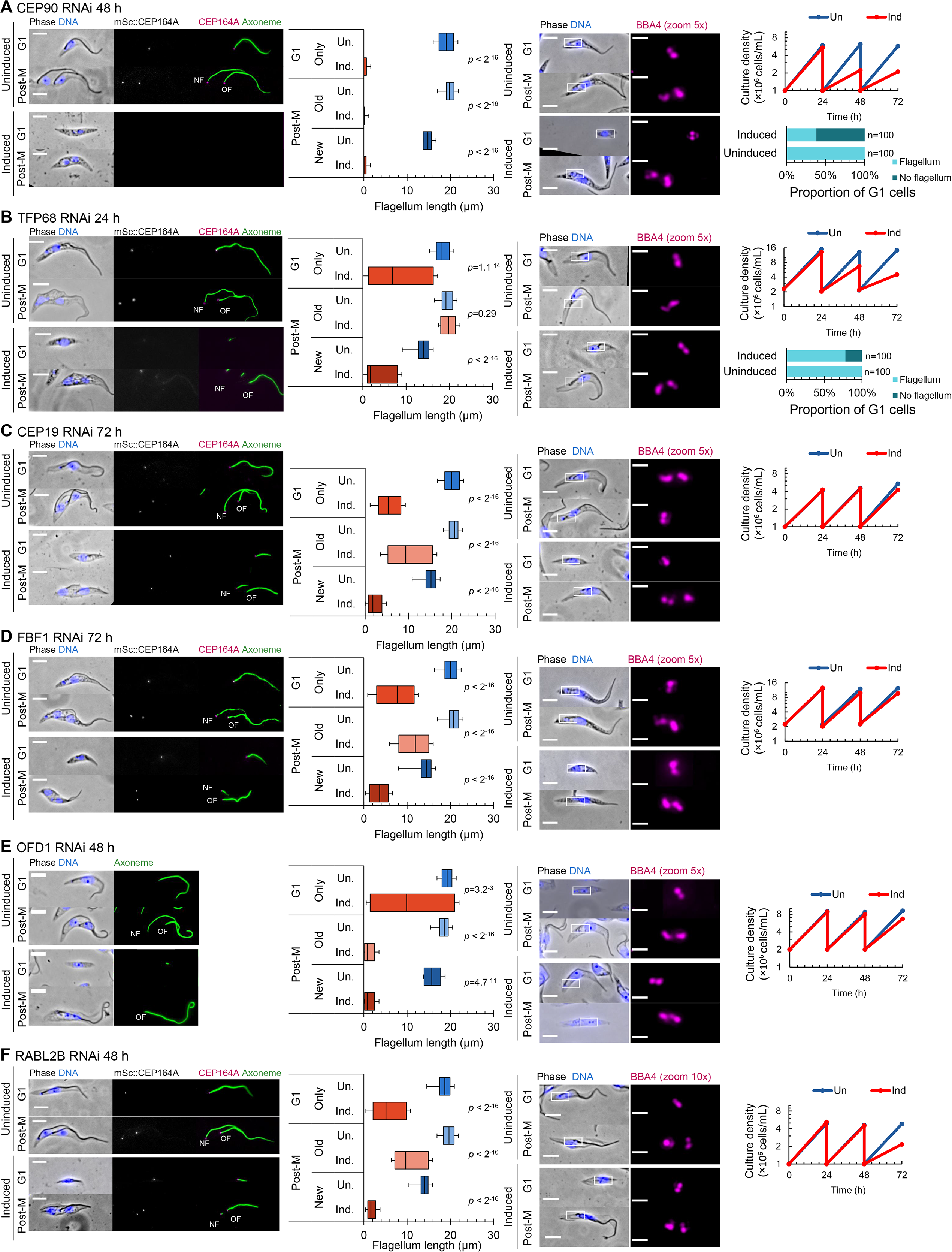
Functional analysis of TFPs essential for new flagellum assembly. RNAi analysis of CEP90 (A), TFP68 (B), CEP19 (C), FBF1 (D), OFD1 (E), RABL2B (F). Panels from left: Representative images of methanol-fixed cytoskeletons with uninduced cells and cells post induction of RNAi. CEP164A endogenously tagged with mScarlet in each RNAi cell line (except for ODF1) is shown in grey/magenta and axoneme (mAb25) in green; Measurements of the axoneme (µm) in G1 cells and the new and old flagella of post mitotic cells in uninduced (Un.) cells and cells following induction of RNAi (Ind.) (n=100); Immunolabelled basal bodies (BBA4) in methanol-fixed cytoskeletons; Growth analysis of cells depleted by RNAi (orange – Ind) and uninduced cells (blue – Un) up to 72 hours; Percentage of cells with no visible flagellum for CEP90 (A) and TFP68 (B). Scale bar = 5µm. NF – new flagellum, OF – old flagellum.

Knockdown by RNAi of CEP19 (Fig 5C), FBF1 (Fig 5D), OFD1 (Fig 5E) and RABL2B (Fig 5F) also resulted in assembly of a very short new flagellum. However, CEP164A remained associated with mature basal bodies in all these cell lines following induction of RNAi knockdown for FBF1, RABL2B or CEP19 (Fig 5C-F), indicating that these proteins are not required for CEP164 localisation, unlike CEP90 and TFP68. We were unable to generate a cell line for OFD1 expressing CEP164A:mScarlet. In all four cell lines (CEP19, FBF1, OFD1, RABL2B), basal body duplication was also unaffected, indicating some level of basal body assembly and maintenance (Fig. 5C-F).

In summary, our data shows that 6 TFPs are essential for new flagellum assembly, but not for basal body biogenesis. Of these only CEP90 and TFP68 were essential for CEP164A recruitment to transition fibres and one of these RABL2B is upregulated in the old flagellum versus the new flagellum in dividing cells. The old flagella remain assembled during induction of RNAi, perhaps suggesting that these TFPs are not essential for flagella maintenance. However, even RNAi knockdown of intraflagellar transport proteins do not perturb assembled flagella length in trypanosomes, suggesting trypanosome flagella are very stable once assembled (Fort et al., 2016).

Next, we asked if RNAi knockdown of TFP122, TFP104 and TFP39 which exhibited a differential localisation pattern between old and new flagella, led to defective new or old flagellum growth, which might suggest a role in regulation of flagellum growth. Cell growth rate was unaffected and CEP164A was retained following RNAi induction. However, measurements of flagellum length did reveal difficulties in reaching the correct flagellum length. For TFP122 both the single flagellum of G1 cells and the new and old flagella of post mitotic cells were significantly longer compared to the uninduced population (Fig 6A). For TFP104 (Fig 6B) the old and new flagella were significantly shorter compared to the uninduced population. There was no cell growth or flagellum length defect in RNAi knockdown of TFP39 (Fig 6C). TFP72 (Fig 6D) also did not reveal any flagellum length defects despite being upregulated in the TF of the new flagellum of dividing cells. In summary TFPs with differential localisation patterns were unable to reach the correct flagellum length. However, TFP49 despite having equal intensity on old and new flagellum also showed a significant reduction in flagellum length, but the cell growth rate was still normal (Fig 6E).

**Figure 6:**
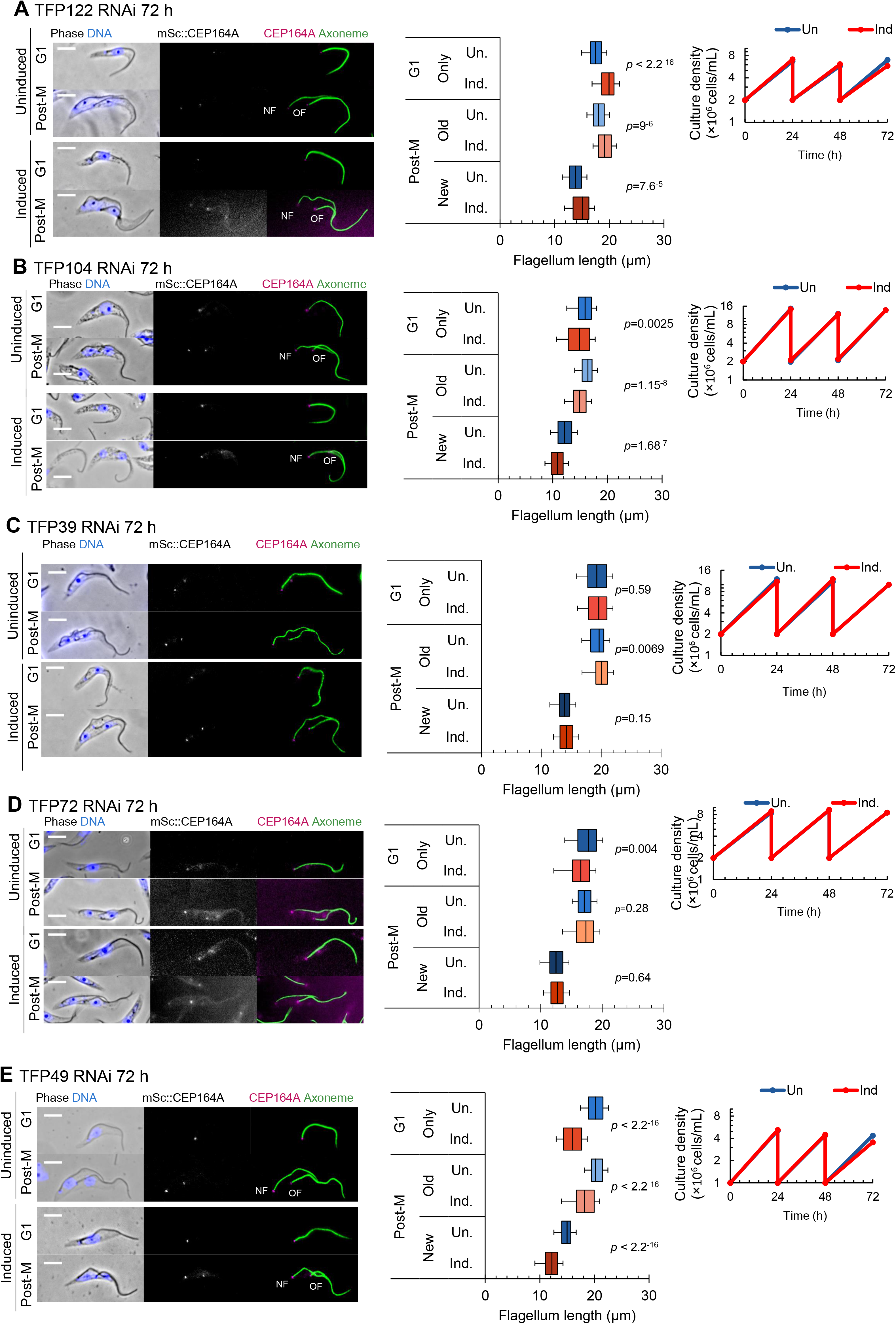
Functional analysis of transition fibre proteins involved in flagellum length regulation. RNAi analysis of TFP122 (A), TFP104 (B), TFP39 (C), TFP72 (D), TFP49 (E). Panels from left: Images of methanol-fixed cytoskeletons with uninduced cells shown above and cells post induction of RNAi shown below. Endogenously tagged CEP164A is in grey/magenta and axoneme (mAb25) in green; Measurements of the axoneme (µm) in G1 cells and the new and old flagella of post mitotic cells in uninduced (Un.) cells and cells following induction of RNAi (Ind.); Growth analysis of cells depleted by RNAi (orange – In) and uninduced cells (blue – Un) up to 72 hours. Scale bar = 5µm. NF – new flagellum, OF – old flagellum.

Finally, depletion of 5 out of the 6 the remaining TFPs chosen for the RNAi knockdown screen (TFP123, TFP17, TFP52, TFP62, SPB1) did not cause a growth defect or flagellum length defect (Supplemental Fig 4A-E). SPB1 was previously shown to localise to the poles of the mitotic spindle during mitosis (Zhou et al., 2018), but here we show that it also localises to the transition fibres during the rest of the cell cycle, highlighting and intriguing link between two microtubule organising centres (Supplemental Fig 1, 4E). Finally, RNAi knockdown of CUL4B (Supplemental Fig 4F), caused a severe growth defect, however, flagellum assembly was not affected and there were no measurable differences in flagellum length between CUL4B depleted and uninduced cells. This data indicates that CUL4B plays a role in cell cycle progression, but might not be involved in flagellum formation or maintenance. Taken together, our functional analysis has identified TFPs essential for ciliogenesis and identified further TFPs that could be important in maintaining correct flagellum length.

## Discussion

### Complexity of transition fibres in ciliated eukaryotes

Our screen identified and confirmed 30 proteins that localise to the TFs in trypanosomes. Two further TFPs were previously experimentally confirmed in Trypanosomes - Retinitis pigmentosa-2 (RP-2) (Stephan et al., 2007) and TFK1 to the matrix between the TF blades (Ramanantsalama et al., 2022) bringing the current number to 32. This challenges the traditional view that TFs are simple structures comprising only a few proteins and demonstrates that trypanosome TFs are complex structures. This is still likely to be an under-estimate as only localisations with a clear mature basal body only signal from the genome-wide protein map (Billington et al., 2023) and where we were able to establish cell lines expressing both the protein of interest and SAS-6 were taken forward in the screen. Given that mammalian and trypanosome TFs are morphologically and structurally similar, we consider it likely that molecular composition of TFs in other organisms has not yet been fully determined and thus speculate there are a number of unidentified TF components in other ciliated eukaryotes. Our automated image analysis interrogated 1000s of individual basal bodies to discern the location and pseudo-diameter for each TFP. The diameter of the TF blades in *T. brucei* observed by STED tomography is ∼338 nm (Trépout et al., 2018), which is similar to our analysis and those using super-resolution microscopy techniques (Tanos et al., 2013; Yang et al., 2018; Bowler et al., 2019; Kumar et al., 2021). Furthermore, our work agrees with these studies in terms of approximate diameter for the human orthologues of CEP164 and FBF1 and with the findings that CEP164 has a wider diameter than FBF1 (Tanos et al., 2013; Yang et al., 2018; Kumar et al., 2021).

To date, C2CD3, OFD1, CEP83, CEP89, SCLT1, CEP164, FBF1, ANKRD26, CEP19, CEP90 and OFD1 are confirmed TFPs in mammalian cells, but not all of these are widely distributed in an evolutionary context, even within Holozoan organisms (Tischer et al., 2021). Our study confirmed 6 mammalian orthologues identified in our screen located to TFs in trypanosomes (CEP164, FBF1, CEP90, OFD1, RABL2B and CEP19). Based on the evolutionary distribution outlined in a bioinformatic study by (Tischer et al., 2021), along with our work, we propose that ODF1, CEP164, FBF1, CEP90, CEP19 and RABL2B may represent a core set of conserved TFPs in eukaryotes. It is likely that *T. brucei* is a highly divergent example of a universal theme in TFP assembly, allowing us to draw some conclusions. Firstly, some *T. brucei* proteins had hints of similarity to cilia/flagella-associated proteins in other eukaryotes based only on predicted structure, suggesting orthology, but sequence divergence could be at the point of detectability by sequence-based methods. Secondly, α helix-rich structural proteins likely tolerate large sequence changes while retaining function, as shown for the transition zone basalin. The α helix-rich transition zone protein Basilin is necessary for axoneme central pair assembly in *T. brucei* and *Leishmania* despite the orthologue having extreme sequence divergence even between these two closely related kinetoplastids – to the point that it was only confidently identifiable due to genome synteny (Dean et al., 2019). It is likely several α helix-rich *T. brucei* TFPs have orthologues in wider eukaryotes that are undetectable by sequence alone.

### Timing of TFP recruitment and ciliogenesis

Trypanosomes have a similar centriole/basal body cell cycle to mammalian cells with pro-basal body duplication also occurring at the G1/S transition. A major difference is in the timing of basal body docking to form a flagellum, which occurs during S-phase immediately before pro-basal body assembly in trypanosomes (Gluenz et al., 2011; Wheeler et al., 2019), unlike mammalian cells where basal body docking and ciliogenesis occurs following cytokinesis. Our work shows that, with the exception of OFD1 and CEP90, TFPs arrive at the newly maturing basal body either immediately before docking or immediately after docking as a new flagellum begins assembly. We identified 15 TFPs arriving at the newly maturing basal body immediately prior to its docking to the flagellar pocket, suggesting roles in membrane docking or initiation of flagellum assembly. Of these 15 proteins only FBF1 exhibited a ciliogenesis defect following knockdown by RNAi, suggesting a high level of redundancy. In contrast, 3 out of the 13 TFPs (TFP68, CEP19, RABL2B) arriving at the maturing basal body immediately after docking exhibited defective ciliogenesis. Both OFD1 andCEP90 were found to arrive on the pro-basal bodies in the previous cell cycle during S-phase, allowing us to conclude that OFD1 and CEP90 are the first TF proteins to be recruited to the pro-basal body prior to maturation in the next cell cycle. Furthermore, depletion of CEP90 and OFD1 by RNAi resulted in a ciliogenesis assembly defect, suggesting roles in TF formation prior to ciliogenesis or initiation of ciliogenesis. Indeed, recent studies demonstrate that CEP90 and OFD1 are involved in building distal appendages in mammals (Kumar et al., 2021). In mammalian cells, TFPs are recruited to the newly maturing basal body at different points during the cell cycle with the appearance of C2CD3 and OFD1 in G2, which is followed in early mitosis by CEP83, CEP89 and SCLT1. Finally, FBF1, CEP164 and ANKRD26 appears in late mitosis with docking occurring following cytokinesis. We can conclude that differences in the timing of TFP recruitment to mature basal bodies in different organisms is likely linked to the differences in the timing of ciliogenesis.

### Transition fibres and ciliogenesis

Several studies have shown that TFPs have essential roles in ciliogenesis and in our work we show that six proteins (CEP90, TFP68, FBF1, CEP19, OFD1 and RABL2B) to be critical for flagellum assembly. Of these, knockdown by RNAi of CEP90 and TFP68 resulted in a loss of CEP164A localisation, suggesting they are also required for correct TF formation. This set of essential proteins include known ciliopathy proteins, already implicated in cilia biogenesis (Focşa et al., 2021). CEP90 is essential for cilium assembly in mammalian cells and forms part of a ciliopathy complex with OFD1 located at the distal end of mother centrioles in mammals (Kumar et al., 2021). In *C. elegans* and mammals FBF1 regulates entry of IFT-A at the ciliary base (Wei et al., 2015), while RABL2B and CEP19 have been shown in mammals to interact with IFT-B complex and function in cilium assembly (Nishijima et al., 2017). Finally, in our study we show that TFP68 is also required for flagellum formation in Trypanosomes and is conserved in other disease-causing kinetoplastida parasites such as *Leishmania spp*. and *Trypanosoma cruzi*, which may represent a kinetoplastid-specific flagellum assembly protein.

In our study we found that basal body duplication was unaffected and basal bodies remained in pairs and located close to the expected region of the cell even in cells where a new flagellum could not be located. We can therefore conclude that TFPs are not required for basal body duplication, but we cannot confirm from our studies if these are necessary for docking *per se*.

### Regulating old flagellum length during new flagellum assembly

A fundamental question in cilia biology is how organisms can faithfully coordinate the assembly of a flagellum to a specific length. In organisms such as *Chlamydomonas reinhardtii,* two flagella are assembled synchronously, where there is equal access to flagellum assembly proteins such as tubulin (Marshall and Rosenbaum, 2001). However, other ciliated/flagellated organisms do not assemble their flagella synchronously, but instead assemble a new flagellum while maintaining the length of an existing flagellum.

Recent studies have implicated a role for TFPs in flagellum length control. CEP164C localises exclusively to the TFs of the old flagellum and functions as part of the old flagellum locking mechanism in trypanosomes (Atkins et al., 2021). Although our screen did not find another TFP where the signal was completely absent on the new flagellum, we did identify four proteins (RABL2B, TFP104, TFP39 and TFP122) with a higher intensity at the base of the old flagellum or new flagellum throughout the cell cycle, suggesting they might be part of this mechanism. Following RNAi knockdown of TFP122, the same phenotype was observed as CEP164C, with cells assembling longer flagella (Atkins et al., 2021). TFP122 contains a kinesin motor head domain and kinesin family proteins have previously been implicated in flagellar length control in trypanosomes (Chan and Ersfeld, 2010). Further studies will be necessary to understand mechanistically their role flagellum length regulation in more detail and how, in general, the TFs ensure fidelity of flagellum length.

We conclude that transition fibres are complex and dynamic structures comprising a core set of evolutionarily conserved components required for assembly and length regulation of eukaryotic cilia and flagella. Our work indicates there may be many additional components to be identified in other ciliated eukaryotes.

## Acknowledgements

We thank The Oxford Brookes Centre for Bioimaging for assistance. We thank Prof. Keith Gull for the kind gift of BBA4 and BB2 antibodies, Prof. Derrick Robinson, University Bordeaux, for kindly gifting the mAb25 antibody and Tryptag.org for access to trypanosome localisation data. Dr Manu Ahmed was funded by a Nigel Groome PhD studentship. The authors declare no competing financial interests.

---

**Supplemental Figure 1: Localisation of 30 endogenously tagged putative transition fibre proteins (TFPs).** TFPs endogenously tagged with SAS-6:mScarlet (magenta). A G1 cell is shown for each tagged TFP. Scale bar = 5µm. M – mature basal body, P – probasal body. Insets represents a 5x zoom. Note, CEP164C is only present on the old flagellum during the cell division cycle and shown here as absent from a G1 cells - see text. Three cell cycle stages shown for SPB1. In G1 cells the signal is only at the transition fibres, during mitosis (M-phase) signal is present on transition fibres and poles of the mitotic spindle, but does not remain post-mitosis when 2 nuclei are visible.

**Supplemental Figure 2: Co-localisation of TFPs with CEP164A or TbRP-2.** TFP cell lines endogenously tagged with CEP164A mScarlet (magenta) or immunolabelled RP2 with the YL1/2 antibody. A G1 cell is shown for each TFP. Scale bar = 5 µm, K – kinetoplast, N - nucleus. Inset represents a 5x zoom.

**Supplemental Figure 3: Measurements of TFs by transmission electron microscopy (TEM);** A, B, C, D, E, F: Longitudinal TEM sections illustrating the organisation of the flagellar pocket membrane (FPM), flagellar axoneme (Ax), mature basal body (MBB), pro-basal body (PBB), densities (D) and transition zone (TZ); B inset: higher magnification TEM of the area in (D) to show the location of transition fibre head densities closely associated with the flagellar pocket membrane. G, H and inset: Cross section TEM of transition fibres; K: cartoon of transition fibre and basal body area and illustration of where the measurement was taken to confirm the width of transition fibres.

**Supplemental Figure 4: RNAi analysis of TFP123 (A), TFP17 (B), TFP72 (C), TFP62 (D), SPB1 (E), CUL4B(F).** Panels from left: Representative images of methanol-fixed cytoskeletons with uninduced cells and cells post induction of RNAi (induced). Endogenously tagged CEP164A in each RNAi cell line (except for TFP17 and SPB1) shown in magenta and axoneme (mAb25) in green; Measurements of the axoneme (µm) in G1 cells and the new and old flagella of post mitotic cells in uninduced (Un) and induced following induction of RNAi (Ind.) (n=100); Growth analysis of cells depleted by RNAi (orange – Ind) and uninduced cells (blue – Un) up to 72 hours; Scale bar = 5µm. NF – new flagellum, OF – old flagellum.

**Supplemental Table 1: TFP general.** Columns from left – Gene ID accession numbers for all TFPs taken forward. Name of each transition fibre protein determined by either orthologue names or molecular weight. Mean pseudo-diameter (nm) (Figure 1E) for each TFP. Mean proximal-distal measurement (nm) (Figure 1D) for each TFP. The timing of recruitment to TF before or after initiation of ciliogenesis.

**Supplemental Table 2: RNAi.** RNAi flagellum length statistical analysis, growth analysis and raw flagellum length data in µm.

## Materials and Methods

### Cell culture

SmOxP9 procyclic trypanosomes derived from TREU 927, expressing T7 RNA polymerase and tetracycline repressor (Poon et al., 2012) were cultured at 28°C in SDM-79 media supplemented with 10% v/v heat-inactivated fetal calf serum (Sigma-Aldrich) (Brun and Schönenberger, 1979). For growth curves, cells were counted every 24 hours using Beckman Coulter V1 cell counter and split to 2 x 10^6^ cells/mL.

### Endogenous cell line generation

N-terminal tagging primers for PCR-only tagging plasmids (pPOT) were designed using http://www.leishgedit.net/Home.html. Constructs for the marker cell line TbSAS-6:mScarlet (Tb927.9.10550) and TbCEP164A:mScarlet (Tb927.5.2440) were generated using the plasmid (pPOTv7) (Dean et al., 2015a). The pPOTv6 plasmid with 3Ty-mNG-3Ty was used to express each putative transition fibre protein. The 51 primer ends are 80 bases of homology allowing homologous recombination into the target locus, when introduced by electroporation (Dean et al., 2015b). Each putative transition fibre protein candidate construct was transfected into the TbSAS-6:mScarlet and/or TbCEP164A:mScarlet cell lines for co-localisation. For transfection, the PCR products were combined with 1x10^7^ cells/mL and transfected using the Amaxa Nucleofactor II system. Transfectants were selected with 20 μg/mL blasticidin or 5µg/mL G418.

### Stable inducible RNA interference (RNAi) cell lines

Primers for RNAi were designed using RNAit (https://dag.compbio.dundee.ac.uk/RNAit/). (Redmond et al., 2003) All constructs for RNAi were generated using the pQuadra stem-loop vector (Inoue et al., 2005). The parental SmOxP9 cell line was transfected with a pQuadra stem-loop plasmid containing the gene of interest and selected with the 10 µg/mL phleomycin (Melford Laboratories).

### Light Microscopy imaging

Light microscopy imaging was carried out using a Zeiss Axio Imager.Z2 wide-field microscope with an ORCA-flash 4.0 Hamamatsu camera (Okerkochen, Germany) and a 100X 1.46NA phase contrast oil immersion objective and all images were acquired using Zeiss Zen blue. For live cell imaging, 1x10^7^ cells were harvested from culture in log phase (5x10^6^-1x10^7^) by centrifugation for 4 minutes at 800 x g. The pellet of cells was washed once in Voorheis modified PBS (vPBS) (137 mM NaCl, 3 mM KCl, 16 mM Na_2_HPO_4_, 3mM KH_2_PO_4_, 46 mM sucrose), resuspended in 1μg/mL Hoescht for 5 minutes and washed twice more and resuspended in vPBS. 1 μL of suspension was settled on an 8-well diagnostic glass slide (Epredia™ - X2XER203B) and mounted with a #0 coverslip (Fisher Scientific – 10011913).

### Fixation methods for light microscopy imaging

Quantitative analysis of pseudo-diameter and distance from SAS-6 measurements were carried out on methanol-fixed cytoskeletons to ensure that the cells are flat to the slide. 1x10^7^ cells were harvested in log phase (5x10^6^-1x10^7^) and centrifuged at 800 x g for 4 minutes. Cells were washed in vPBS twice and resuspended in vPBS before settling onto slides. Cytoskeleton extract was carried out using 1% IGEPAL-NP40 (Sigma-Aldrich - I8896) in PEME buffer (100 mM PIPES-NaOH pH 6.9, 1 mM MgSO_4_, 2 mM EGTA, 0.1 mM EDTA) before incubation in methanol and left for 30 minutes at -20°C. Slides were rehydrated in PBS at room temperature for 5 minutes, stained with 1 μg/mL Hoescht 33342. Formaldehyde-methanol fixation was used for IFT172 primary antibody labelling. Cells were harvested in log phase (5x10^6^-1x10^7^/ml) by centrifugation at 800 x g for 4 minutes. The pelleted cells were washed twice (vPBS), settled onto glass slides and left to settle for 10 minutes before being treated for 5 minutes with formaldehyde using 16% methanol free formaldehyde (Thermo Scientific™ 28908) in PBS. Slides then incubated in methanol at -20°C for 5 minutes and were subsequently rehydrated in PBS for 5 minutes at room temperature before being processed for immunofluorescence.

### Expansion microscopy (ExM)

Cells were harvested and pelleted by centrifugation for 4 minutes at 800 x g and washed twice in vPBS before being resuspended in fixation solution consisting of 4% formaldehyde and 4% acrylamide and transferred to a 12 mm round coverslip (Sigma, P4707) coated in poly-L-lysine and placed in a 24 well plate. Cells were fixed overnight at room temperature.

For gelation 90µl monomer solution (19% sodium acrylate (Sigma, 408220), 10% acrylamide and 0.1% N, N’- Methylenebisacrylamide (Sigma, M7256) in PBS) was combined with 5 µL of 5% N,N,N’,N’-tetramethylethylenediamine (TEMED) (Sigma, T9281) and 5 µL of 5% ammonium persulfate (APS) (Thermo Scientific, 17874). 50 µl was added to a strip of parafilm in a humidity chamber on ice. Coverslips (cells faced down) were placed on top and incubated for 5 minutes, transferred to 37°C and incubated for a further 30 minutes. Gels were detached using a few drops of ultrapure water Specimens were transferred to denaturation buffer (50mM tris(hydroxymethyl)aminomethane-hydrochloride (Sigma, 77-86-1), 200 mM sodium chloride (Sigma, S9888) and 200mM sodium dodecyl sulphate (FisherSci, 151-21-3), pH 9) in a 6-well plate. Gels were transferred to a microcentrifuge tube containing 1 mL denaturation buffer and denatured at 95°C for 90 minutes. Denaturation was followed by 3x20 minute expansions in ultrapure water in a 10 cm dish. A 10 x 10 mm section of gel was cut for antibody staining. Antibodies were diluted in 2% BSA and gels incubated with 500 µL of primary antibody overnight, 3 x 20 minute washed in ultrapure water, the secondary antibody incubation overnight followed by 3 x 20 minute washes in ultrapure water.

Specimens were finally incubated with 10 µg/mL Hoechst (Thermo Scientific, 622249) in PBS for 30 minutes. Cells were imaged on a 35 mm glass bottom imaging dish (Thermo Scientific, 150680) using a Zeiss 880 confocal with AIRYSCAN.

### Antibodies and Immunofluorescence

Slides pre-blocked with 1% bovine serum albumin (BSA) in PBS for 30 minutes. Primary antibodies were diluted in PBS containing 1% BSA in a humidity chamber for 1 hour at room temperature and washed 5 x 5 minutes in PBS. Secondary antibodies were diluted in PBS containing 1% BSA left in a humidity chamber at room temperature for 1 hour. Flouresein (FITC) AffiniPure F(ab’)2 fragment Donkey anti-Mouse IgG (H+L) (Jackson Immunoresearch), Rhodamine Goat Anti-Mouse IgM for BBA4 primary (Jackson Immunoresearch). Slides were washed 5 x 5 mins with PBS and 1 μg/mL Hoescht.

### Transmission electron microscopy

25% glutaraldehyde (TAAB) was added to the culture suspension to a final concentration of 2.5%. Cells were pelleted and resuspended in a primary fixative containing 2.5% glutaraldehyde, 2% paraformaldehyde (Agar Scientific), and 0.1% tannic acid (TAAB) in 0.1 M phosphate buffer, pH 7.0 (Sigma-Aldrich). Cells were fixed for 2 h at room temperature. Pellets were washed with 0.1 M phosphate buffer (pH 7.0) and postfixed in 1% osmium tetroxide (Agar Scientific) in 0.1 M phosphate buffer (pH 7.0) for 1 h at room temperature. Samples were rinsed and stained en bloc for 40 min in 2% aqueous uranyl acetate (TAAB), dehydrated in an ascending acetone series (Fisher Scientific), and embedded in hard formulation Agar 100 resin (Agar Scientific). 70-nm thin sections were made and stained using lead citrate for 5 min, followed by three washes with MilliQ water. Images were captured using a Hitachi H-7650 transmission electron microscope. Method as per (Atkins et al., 2021).

### Ortholog identification

To identify orthologs of *T. brucei* TFPs, we used a primarily reciprocal best protein BLAST search (version 2.9.0), accepting reciprocal hits with e ≤ 10^-5^. supplemented with orthogroup detection using Orthofinder (version 2.5.4, using FastME 2.1.4) (Emms and Kelly, 2015, 2019). We identified orthologs in a diverse set of eukaryotes with previously-described ability to form basal bodies and distal appendages (Carvalho-Santos et al., 2011b). Predicted proteome sequences were primarily from NCBI genomes: GCA_001641455.1_Mp_v4 (*Marchantia polymorpha*), GCA_023343905.1_cvul (*Chlorella vulgaris*), GCF_000002825.2_ASM282v1 (*Trichomonas vaginalis*), GCF_000004985.1_V1.0 (*Naegleria gruberi*), GCF_000004695.1_dicty_2.7 (*Dictyostelium discoideum*), GCF_000091205.1_ASM9120v1 (*Cyanidioschyzon merolae*), GCF_000002595.1_v3.0 (*Chlamydomonas reinhardtii*), GCF_000214015.3_version_140606 (*Ostreococcus tauri*), GCF_000001735.4_TAIR10.1 (*Arabidopsis thaliana*), GCF_000002865.3_V1.0 (*Monosiga brevicollis*), GCF_000002985.6_WBcel235 (*Caenorhabditis elegans*), GCF_000001215.4_Release_6_plus_ISO1_MT (*Drosophila melanogaster*), GCF_000209225.1_ASM20922v1 (*Nematostella vectensis*), GCF_000203795.1_v1.0 (*Batrachochytrium dendrobatidis*), GCF_000328475.2_Umaydis521_2.0 (*Ustilago maydis*), GCF_000149755.1_P.sojae_V3.0 (*Phytophthora sojae*), GCF_000189635.1_JCVI-TTA1-2.2 (*Tetrahymena thermophila*) & GCF_000002765.5_GCA_000002765 (*Plasmodium falciparum*). Trypanosomatid parasite predicted proteome sequences were from TriTrypDB.org version 59, part of VeuPathDB (Amos et al., 2022): TbruceiTREU927 (*Trypanosoma brucei*), TcruziDm28c2014 (*Trypanosoma cruzi*) & LmexicanaMHOMGT2001U1103 (*Leishmania mexicana*). Human and yeast predicted proteome sequences were from UniProt *Homo sapiens* (UP000005640) and *Saccharomyces cerevisiae* (S288C) genome database respectively.

### Protein domains and Alphafold

Protein domains were identified using the *T. brucei* genome database at TriTrypDB.org, part of VeuPathDB(Amos et al., 2022). Protein structures were predicted using AlphaFold2 (Jumper et al., 2021a) using a previously-described custom sequence database to generate the input multiple sequence alignments (Wheeler, 2021), to provide better sequence diversity of the Discoba lineage in which *T. brucei* sits. Predicted protein secondary structure was extracted from the three-dimensional predicted structure using DSSP(Kabsch and Sander, 1983), summarising structure term “H” for α-helices and “E” or “B” as β-sheets. Protein structures were visualised using PyMOL. Protein structure searches against PDB were carried out using the FoldSeek server(Kempen et al., 2022)

### Automated Image Analysis for measurements of distance and pseudo-diameter

An ImageJ/FIJI macro was developed for automated analysis of distance of the centre of signal between each tagged cell line co-expressed between each TFP::mNG and SAS-6::mSc TFP to determine the dimensions of the tagged TFP structure. SAS-6 and tagged TFP protein signal foci in the green and red fluorescent images were first either selected manually using the ImageJ/FIJI point selection tool or identified automatically using the "Find Maxima" function in ImageJ/FIJI. The precise centre points were determined using a Gaussian fit of signal in the x and y directions for each point in the green and red fluorescent images separately (as previously described https://link.springer.com/protocol/10.1007/978-1-0716-0294-2_24). TFP::mNG and SAS-6::mSc foci lie very close (within ∼500 nm), within the same cell, so distance from each focus to the nearest SAS-6 focus, excluding distances > 500nm, was taken as the distance between these structures. To determine the dimensions of the TFP structure, the centre point of the mNG-tagged TFP structure in the green fluorescence image was identified as above. Next, signal intensity across the TFP structure was sampled in 45 straight lines spanning across the centre point, with the lines positioned at 0 degrees (horizontal) through to 180 degrees at 4 degree steps. For each line, a Gaussian was fitted and the standard deviation was taken as an effective diameter along that line. The largest effective diameter was taken as the TFP structure pseudo-diameter, excluding outliers >∼500 nm.

### Flagellum measurements and statistical analysis

Measurements of the axoneme were performed using FIJI software. G1 cells (a single nucleus) and post mitotic cells (2 separate nuclei) were identified by Hoechst staining of the DNA. immunolabelled axoneme (Mab25 – see above) was traced using a segmented line and a curve was added using the ‘fit spline’ function on FIJI. Measurements were recorded and transferred to a Microsoft excel spreadsheet. Boxplots representing flagellum lengths were generated using the Microsoft excel. Statistical analysis of flagellum length (Mann-Whitney U test) was performed in R studio.

### Quantification of image intensity

All measurements of intensity of fluorescent signal were made in FIJI. For quantification of transition fibre signal intensity between basal bodies in dividing cells, a rectangular area of interest of a defined size was generated and the integrated density was measured within the box. Measurements were made at the transition fibres and around the posterior end of the cell to measure the background signal. The background signal was subtracted from the transition fibre signal at the base of the new flagellum and old flagellum (n=50).

