## Supplementary figures and images for "Identification of 30 transition fibre proteins reveals a complex and dynamic structure with essential roles in ciliogenesis"

### supplemental figure 1

#### Supplemental Figure 1

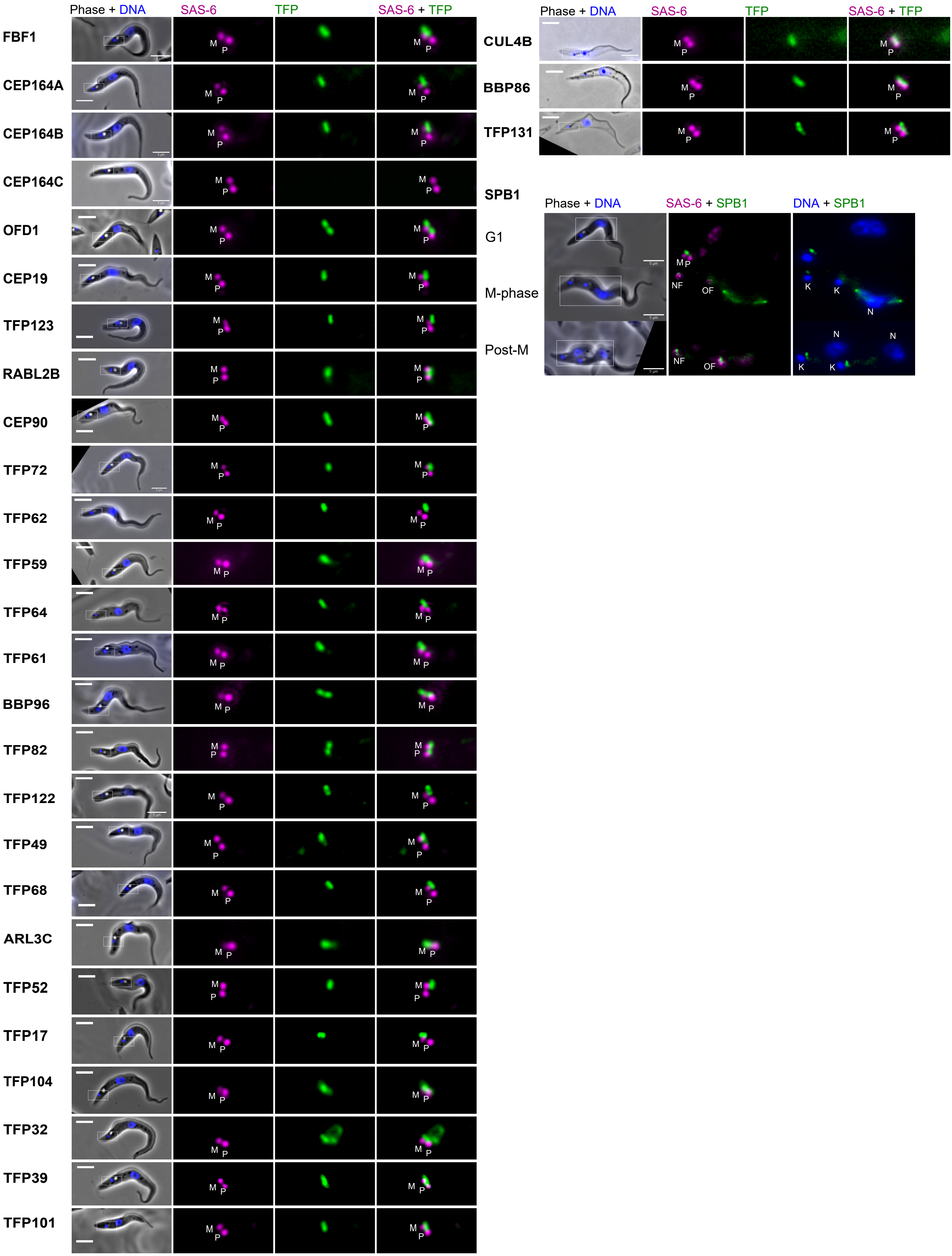

### Supplemental figure 2

#### Supplemental Figure 2

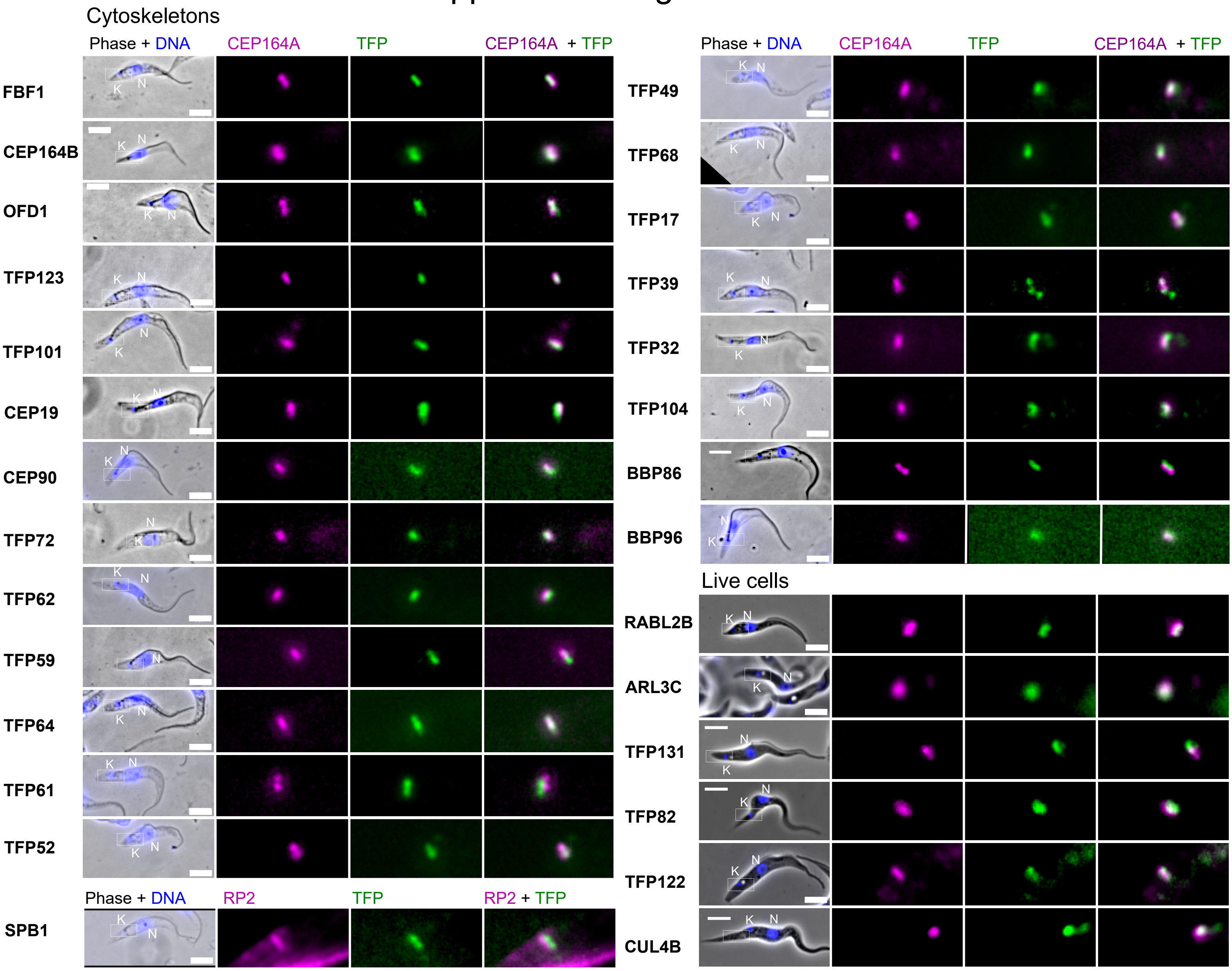

### Supplemental figure 3

#### Supplemental Figure 3

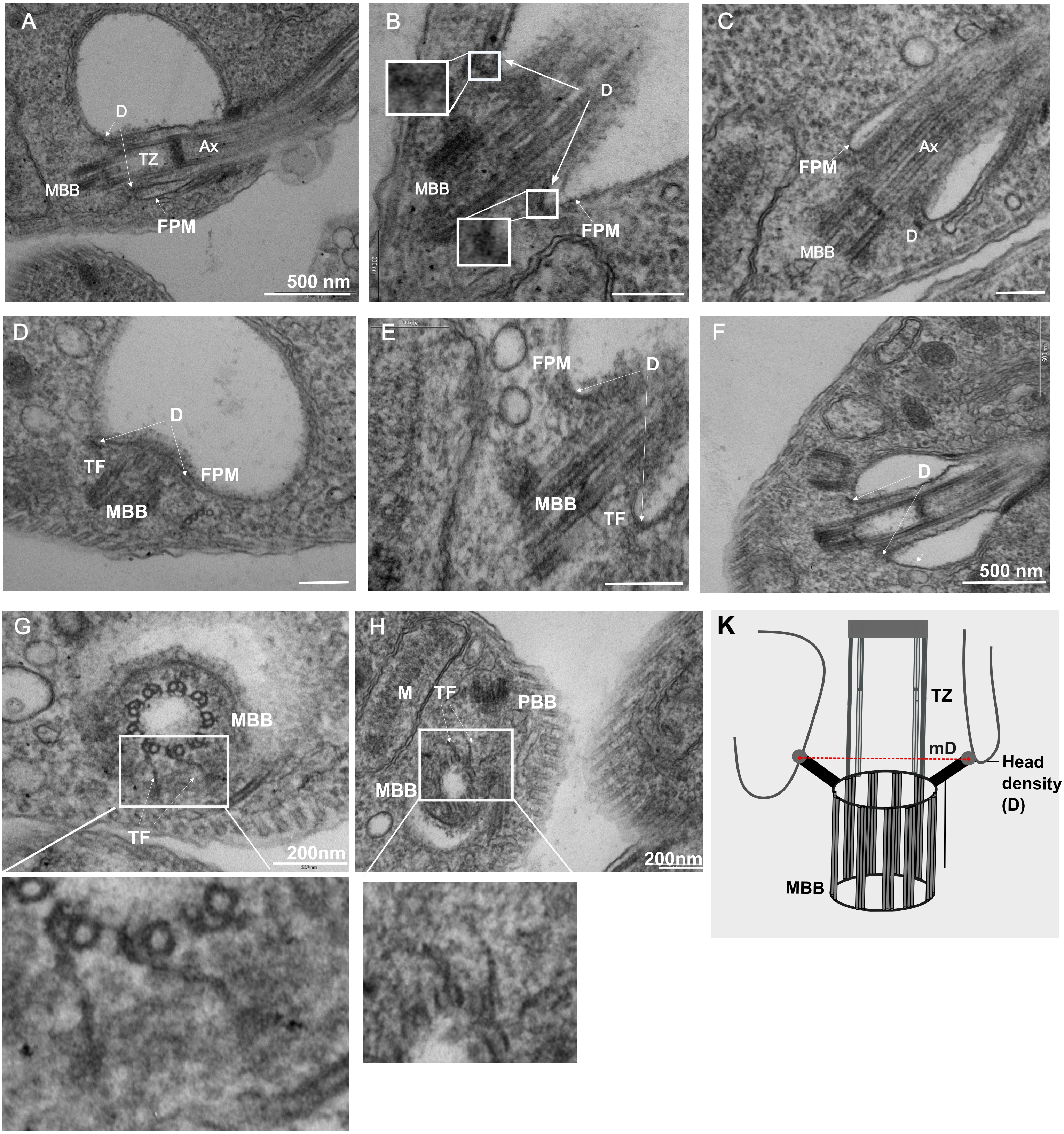

### Supplemental figure 4

#### Supplemental Figure 4

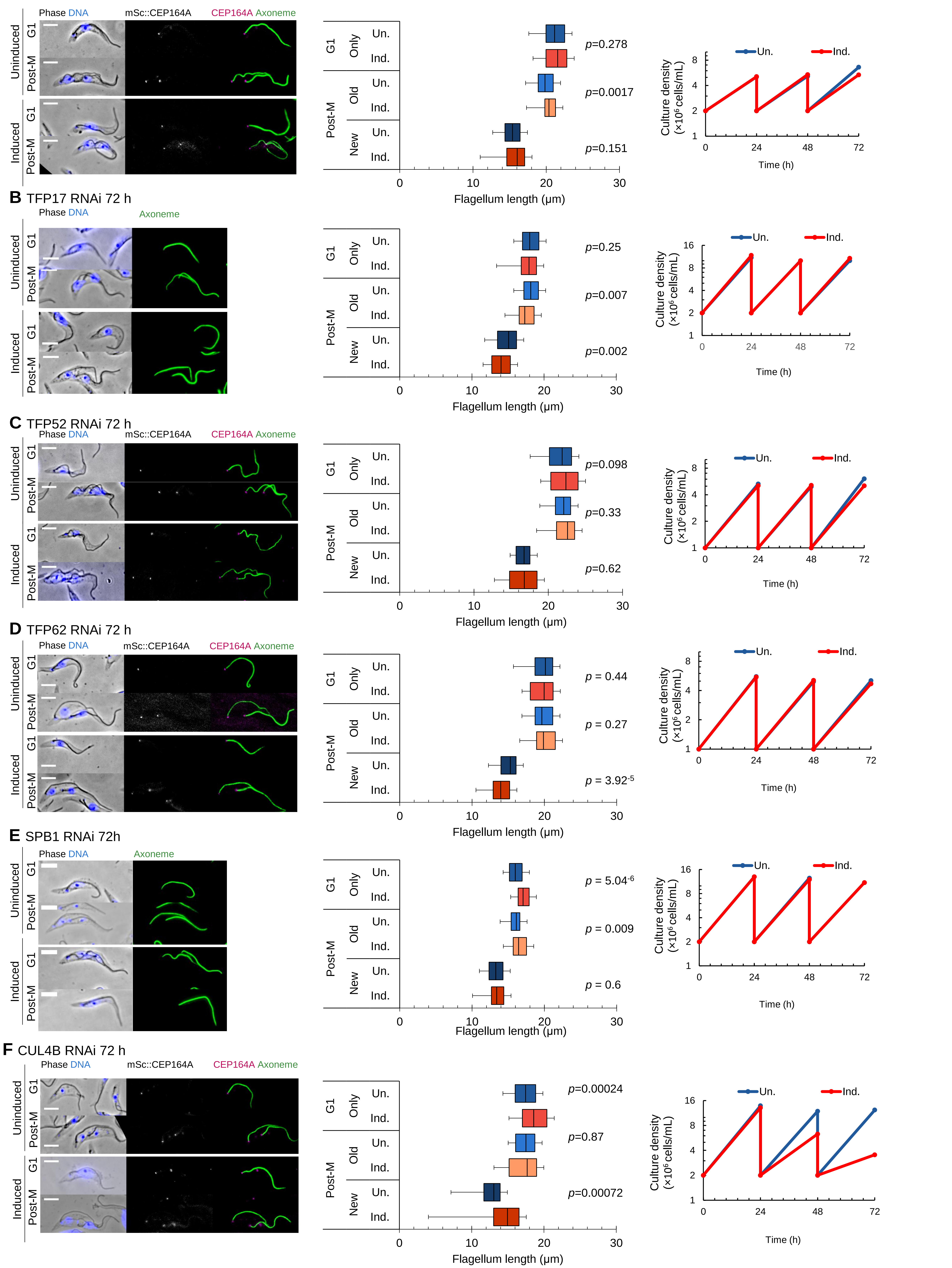
